# A control engineering perspective on the advantages of efference copies

**DOI:** 10.1101/2023.07.24.550357

**Authors:** Benjamin P. Campbell, Huai-Ti Lin, Holger G. Krapp

## Abstract

Biological systems have evolved to perform high-speed voluntary movements whilst maintaining robustness and stability. This paper examines a control architecture based on the principles of efference copies found in insect sensorimotor control which we call the fully-separable-degrees-of-freedom (FSDoF) controller. Within a control engineering framework, we benchmark the advantages of this control architecture against two common engineering control schemes: a pure feedback (PFB) controller and a Smith predictor (SP). Our study identifies three advantages of the FSDoF for biology. It is advantageous in controlling systems with sensor delays, and it can effectively handle noise. Thirdly, it allows biological sensors to increase their operating range. We evaluate the robustness of the FSDoF controller and show that it achieves improved performance with equal stability margins and robustness. Finally, we discuss variations of the FSDoF which theoretically provide the same performance.

## 1 Introduction

Biological systems use a different architecture to control stabilization reflexes and goal-oriented behaviours compared with those frequently used in engineering. A typical feedback controller (Fig. 1a), works by sensing the current state, e.g. position, velocity, angle, and subtracting that from the desired state to give an error signal. The error is provided to a controller which generates a correcting actuation command, e.g. force, voltage, torque. The actuation acts on the dynamics of the system to reduce the error, thereby attempting to maintain equal current and desired states [1].

**Figure 1:**
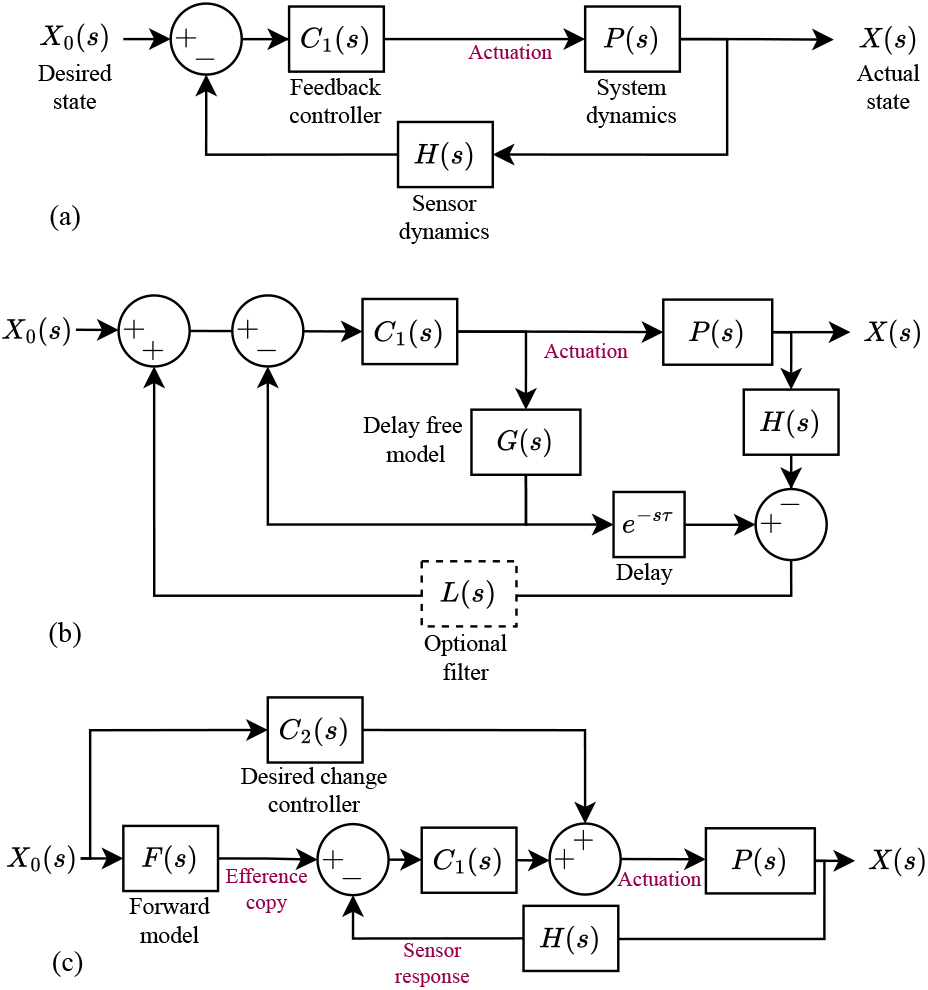
Three benchmarked control architectures. Blocks include Laplace transforms of the respective dynamics. Red labels name the signals between blocks. Signals labelled “actuation” refers to actuation commands, the actuator dynamics are included in *P*(*s*). (a) Pure feedback controller (PFB): the state is measured by sensors with some dynamics, this is subtracted from the desired state to give an error. The feedback controller, *C*_1_(*s*), generates an actuation command from the error. (b) Smith predictor (SP): a feedback controller, *C*_1_(*s*), controls an undelayed plant model and sensor dynamics (*G*(*s*)), to generate the command for the real system *P*(*s*). (c) FSDoF controller: the feed-forward, desired change controller, *C*_2_(*s*), generates the motor command to actuate the desired state. The desired state is also used by a forward model to predict the sensory consequences and generate an efference copy. That efference copy signal cancels the expected sensor response. The feedback controller, *C*_1_(*s*), generates an output only if the sensor response and efference copy do not match.

Previously it was suggested that the sensorimotor control in insects differs by having two layers: an inner-loop and an outer-loop [2, 3, 4]. The inner-loop consists of a stabilising controller that compensates for any perceived change in state [5]. Inner-loop control alone would result in a system that is unable to change state [4]. The outer-loop then has two roles which accommodate desired changes in state. Firstly, it provides an actuation command to the motor system to carry out the desired state change. Secondly, it cancels the expected sensor response resulting from the actuation command. This, in turn, prevents the stabilising controller in the inner-loop from counteracting the desired state change (see [4]). In biology, this expected sensor response is called an efference copy [6, 7, 8, 9, 10].

In recent years, some experimental studies have provided physiological evidence for efference copies in insects. Combining neurophysiological and behavioural methods Kim et al. (2015) found that insects modify their sensor response during spontaneous yaw rotations using efference copies [11]. It was then shown that the cancellation signal scaled across different cells with the optic flow each cell should expect [12]. Later, Fenk et al. showed that this was not the case for actuation in response to external disturbances [5]. In addition, experiments on the speed control in *Drosophila* have shown that octopamine increases the gain of optic flow processing interneurons. Yet, when octopaminergic neurons are silenced, there is no change in average flight speed [13]. The physiological [11, 12, 5, 14] and behavioural [13] evidence strongly support and align with, the inner-loop and outer-loop structure for insect sensorimotor control.

A control architecture that uses efference copies, predicting sensor responses to desired changes in state, is not unique to insects; it is also found in primates [7]. However, compared to insects cellular evidence for its implementation is more challenging to obtain in primates.

Here we use the term forward model in the way it is often applied in the context of insect sensorimotor control [15]. It is a function, or filter, that maps a desired change in state into an expected sensor response. It is not an internal map of the world or an understanding of what will happen when certain decisions are taken, like what is described in [16]. It is also different to the concept of a forward model in much of the cerebellum literature [17, 18, 19, 20, 21], where typically a forward model refers to a model of the dynamics of the system. Whilst there are many hypotheses for different cerebellum control schemes, the most typical uses a model of the dynamics of the system to find an inverse model which can be used to generate a motor command [17]. Here, we refer to a forward model as a filter that predicts a sensor response from a desired state change. It is therefore not just a model of the dynamics, but a filter that models the insect controller, the dynamics, and the sensor properties.

The second aspect of the outer-loop is the generation of the actuation command to execute the desired state change or desired movement. This feedforward controller is another function, or filter, that this time maps a desired state change into a motor command resulting in muscle contraction. Mostly in engineering control theory feedforward controllers are used to supplement the actuation of the feedback controller when changing desired state, referred to as a two-degree-of-freedom (2DoF) controller [22]. We differentiate this type of feedforward control from what is hypothesised in the insect sensorimotor system by name and by a simple principle. We refer to the feedforward controller in the insect sensorimotor system as a desired change controller; where the principle difference is that instead of supplementing the actuation of a feedback controller, it provides *all* of the actuation commands needed to change state.

Considering their different tasks: the inner-loop enabling compensatory motor action in response to external perturbations and the outer-loop enacting goal-directed behaviours, the architecture was, so far, referred to as an inner-outer loop controller. To avoid confusion with other control architectures we will refer to it as a fully-separable-degrees-of-freedom-controller (FSDoF); it is a specific formulation of a 2DoF controller [22] where the degrees of freedom are completely separated. In the first half of this paper, we compare the performance of the FSDoF controller and two common controllers in engineering to identify its advantages. Then, we discuss the robustness of the FSDoF control scheme. Finally, considering the design constraints of biological systems, we address the question of why the FSDoF controller appears to be employed as opposed to other structures with similar advantages. The three controllers compared are given in Figure 1. The feedback controller (Fig. 1a), which we will refer to as the *pure* feedback controller (PFB), to differentiate it from the feedback component of the other two controllers. The Smith predictor (SP) (Fig. 1b), and the efference copy inspired fully-separable-degrees-of-freedom (FSDoF) controller (Fig. 1c), combining the desired change controller, forward model, and feedback control.

This paper is the first to analyse the biology-inspired FSDoF control system and compare it with other typical controllers. The neural control structure has evolved over hundreds of millions of years enabling the integration of stabilization reflexes and goal-oriented behaviours under severe energy constraints. By relating a control systems analysis of the structure, to the common sensor and dynamic properties of insects, we attempt to answer: what advantages does biology gain from efference copy control methods?

## 2 Formulation of the FSDoF insect sensorimotor control architecture

By far the most widespread and convenient way to examine control architectures is using the complex frequency (Laplace) domain [1]. It is a mathematical tool that allows interactions between temporal filters to be treated algebraically, and convolution to be treated as multiplication.

There are three main metrics used to assess the suitability of a controller: (i) disturbance rejection, how quickly can the controller drive the actuators to remove external perturbations (e.g. due to gusts of wind), (ii) performance in reference tracking, i.e. how well the actual state tracks the desired state, and (iii) robustness, i.e. the stability of the system if the controller is built around an incorrect systems dynamics model.

By finding the transfer functions of the closed-loop controlled systems we can quantify these metrics. First, looking at disturbance rejection: the transfer function from a disturbance *U*(*s*) to the actual state *X*(*s*), is given by equation 1. Where *P*_*d*_(*s*) is the transfer function for how the disturbance affects the state.

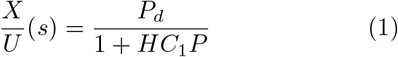

The same equations are obtained for all three architectures in Figure 1. If all three architectures use the same feedback controller, they will all have the same performance in handling disturbances. This is intuitive, since in order to reject a disturbance it must be sensed, and therefore feedback control must be used to remove its impact.

For performance in tracking the desired state, all three controllers have different transfer functions. The transfer functions for the pure feedback controller (PFB, Fig. 1a) and SP (Fig. 1b) are given in equations 2 and 3 respectively.

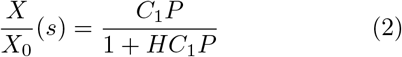

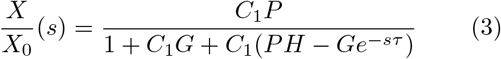

However, for the SP, *G*(*s*) is an undelayed model of the undelayed system and sensor dynamics (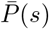 and 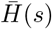, respectively). Therefore, we assume 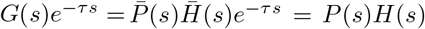, i.e. the estimated model of the system is perfect. We will break this assumption when we discuss robustness to modelling errors in Section 4.4. For now, the resulting transfer function is given by equation 4.

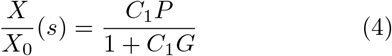

The only difference between the two is that the PFB controller is designed around the system and sensor dynamics including the delay, and the SP controller is designed around a delay-free system.

The FSDoF control architecture (Fig. 1c) has the transfer function given by equation 5.

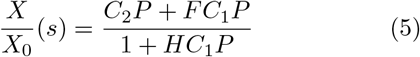

However, the forward model *F*(*s*) is not a random filter, it should turn the desired state changes into the expected sensor response. By looking at Figure 1c, following the loop from desired state to sensor response, we can see that the forward model should predict:

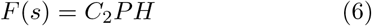

Substituting this into equation 5 we get a different-looking transfer function from desired state to actual state given in equation 7.

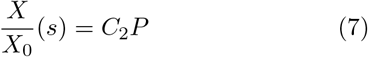

This effectively corresponds to open-loop control. Because there is a feedback loop, however, the disadvantages of *pure* open-loop control do not apply to the FSDoF architecture. The architecture can still compensate for external disturbances, and it can still deal with unstable dynamics.

For the first control metric (i) disturbance rejection, the equations show that all three controllers will have the exact same performance, given the same *C*_1_(*s*). This is not the case for the second control metric (ii) reference tracking, where the three architectures are formalised by different equations; the majority of the rest of the paper will be on simulations showing the differences in performance. Then finally (iii) the robustness of the controller to differences in the real and expected system dynamics will be discussed in Section 4.4.

## 3 Methods

To explore the differences in reference tracking performance between the efference copy-inspired FSDoF controller and the two typical engineering controllers we follow a simple procedure: select some system and sensor dynamics, design all three controllers around controlling those dynamics, and compare the response to a step change in the desired state. The responses of the closed-loop systems were simulated using a discrete approximation (ode45) of continuous time in Simulink MATLAB.

The controller architectures were tested on 36 different system dynamics. Table 1 gives the structures, chosen to represent different common dynamics: 1 and 3 represent low pass filter dynamics, 2 includes an integrator, 4 and 6 include a *zero*, and 5 is an unstable system. Each of the six plant models was parameterised with different bandwidths and sensor delays, *ω* and *τ* respectively. This was to provide a range where either the sensor delay or the apparent delay due to the system dynamics dominated. We used three different sensor delays: 0 ms, 50 ms, and 100 ms. Each with two bandwidths: 10 rad s^*−*1^, and 50 rad s^*−*1^. This range was selected to approximately reflect the range of band-widths and delays in the insect control system from system identification results, where the animal is enacting a desired behaviour [23, 24]. The frequency of the *zero*(*ω*_*z*_) was set to be 30 rad s^*−*1^, between 10 rad s^*−*1^ and 50 rad s^*−*1^ meaning the pole would be both dominant and non-dominant. To test each controller on the 36 dynamics, we had 84 different controllers: 36 different PFB and FSDoF controllers, and 12 different SPs since the SP is designed around the undelayed system.

**Table 1:**
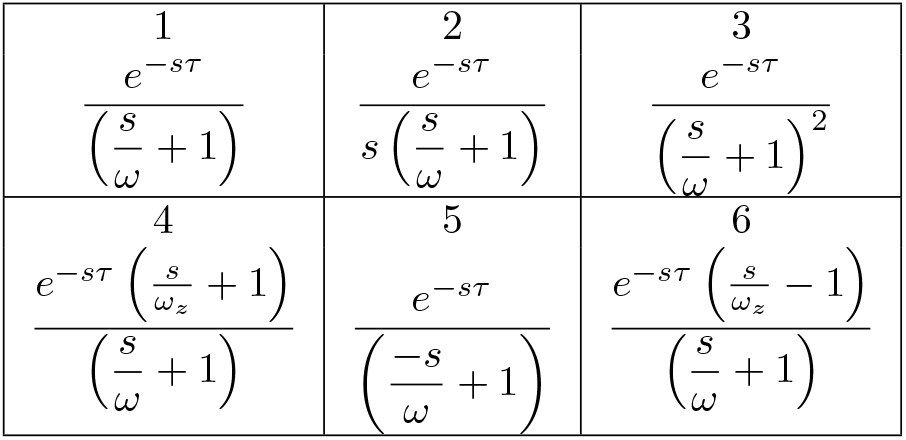
Six different combinations of transfer functions describing system and sensor dynamics, *P*(*s*)*H*(*s*), as shown in Figure 1.

For a fair comparison, the limits of the actuators for each of the three systems must be approximately the same. To ensure this, the maximum allowable actuation of the FSDoF control architecture was not allowed to exceed 1% of the maximum actuation used by the PFB controller. The feedback controller was a proportional-integral-derivative (PID) controller. For the PFB and the SP, the feedback controller was configured using MATLAB’s optimal PID tuner with a design focus on reference-tracking, and a desired phase margin of 45^*°*^. This meant there were three optimised parameters (gains) dictating the reference tracking performance for the SP and PFB. The FSDoF inherently has more filters and parameters. To limit the degrees of freedom optimising the FSDoF a predefined structure for *F*(*s*) and *C*_2_(*s*) was determined. Subsequently, reference tracking performance was tuned with a single bandwidth parameter. Thus, the optimisation process involved three variable parameters for the SP and PFB and one for the FSDoF (Supp. Sec. 2). When the MATLAB PID tuner was unable to find a stabilising controller that met the requirements, that case was omitted from the analysis.

An example step response is given in Figure 2. The MATLAB optimal gains for the PFB controller were *K*_*p*_ = 0.86 and *K*_*i*_ = 5.66, for the SP controller the feedback gains were *K*_*p*_ = 1.72 and *K*_*i*_ = 19.59. The feedback gains in the FSDoF do not impact the reference tracking behaviour. However, to obtain the results in Figure 2, a feedback controller was tuned with a design focus on disturbance rejection which had gains of *K*_*p*_ = 0.70 and *K*_*i*_ = 6.28. More relevant to the reference tracking is *C*_2_(*s*) which was chosen to have a *zero* at 10 rad s^*−*1^, and have two poles at 29 rad s^*−*1^. Correspondingly, the forward model became a second-order low-pass filter with the same poles. The data on time delays was collected by repeating this process for all 36 different systems.

**Figure 2:**
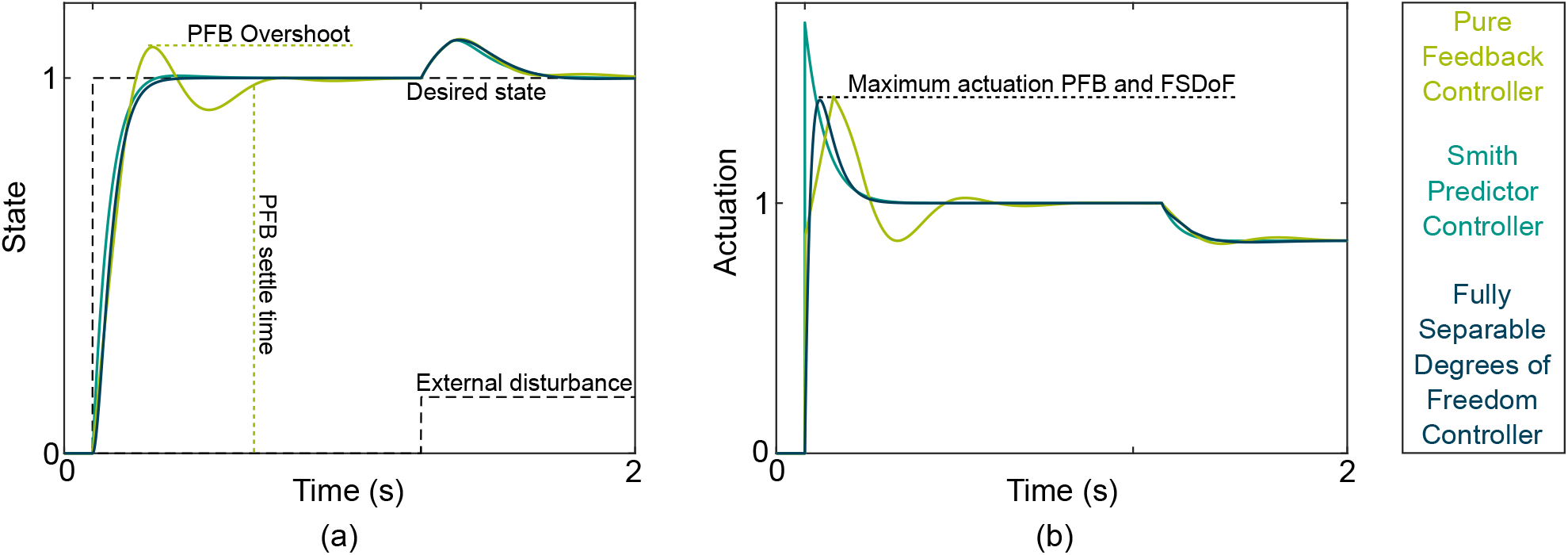
Step response of all three controllers on the first structure of dynamics in Table 1 with *ω* = 10 rad s^*−*1^ and *τ* = 100 ms. (a) Desired state (dashed step function) and the actual state, the PFB overshoots 8.3 %, the SP overshoots 0.6 %, whilst the FSDoF controller does not overshoot. The FSDoF controller has a settle time of 20 ms, compared with 56 ms for the PFB and 18 ms for the SP controller. (b) shows the actuation used by the three controllers. The maximum actuation for the FSDoF and PFB is 1.4 units, and for the SP it is 1.7.

The demonstration of handling noisy systems was done by tuning two PFB controllers to the system dynamics and only altering the response time parameter in the PID tuner, and therefore the bandwidth. It is not possible to perfectly simulate white noise due to its infinite frequency spectrum. Instead, a white noise approximation using a random sequence with a correlation time constant was injected at the sensor. The correlation time constant must be significantly smaller than the smallest time constant in the system being simulated, therefore, we used a value of 0.1 ms and a power of 0.001. The FSDoF controller was given the same feedback controller as the lower bandwidth PFB. The constraint of maximum actuation was relaxed for this demonstration, given the actuation used was largely determined by the response to noise.

## 4 Results

An example of how the three controllers perform when designed around the same system dynamics is given in Figure 2. To assess the performance of the reference tracking we measured the overshoot and the settle time. The overshoot measures how far the actual state exceeds the desired state before converging to the desired reference state; and the settle time is the time to reach and stay within 2 % of the desired state as shown in Figure 2a. We also measured the actuation costs in the form of the maximum actuation required as shown in Figure 2b.

### 4.1 Advantage 1: Mitigating the impact of sensor delays

The performance results for the overshoot and settle time at different sensor delays are shown in Figure 3. The fully-separable-degrees-of-freedom (FSDoF) controller never overshoots by design (Fig. 3b). The other two controllers do overshoot, and in the case of the pure feedback controller (PFB), the overshoot increases with the sensor delay.

**Figure 3:**
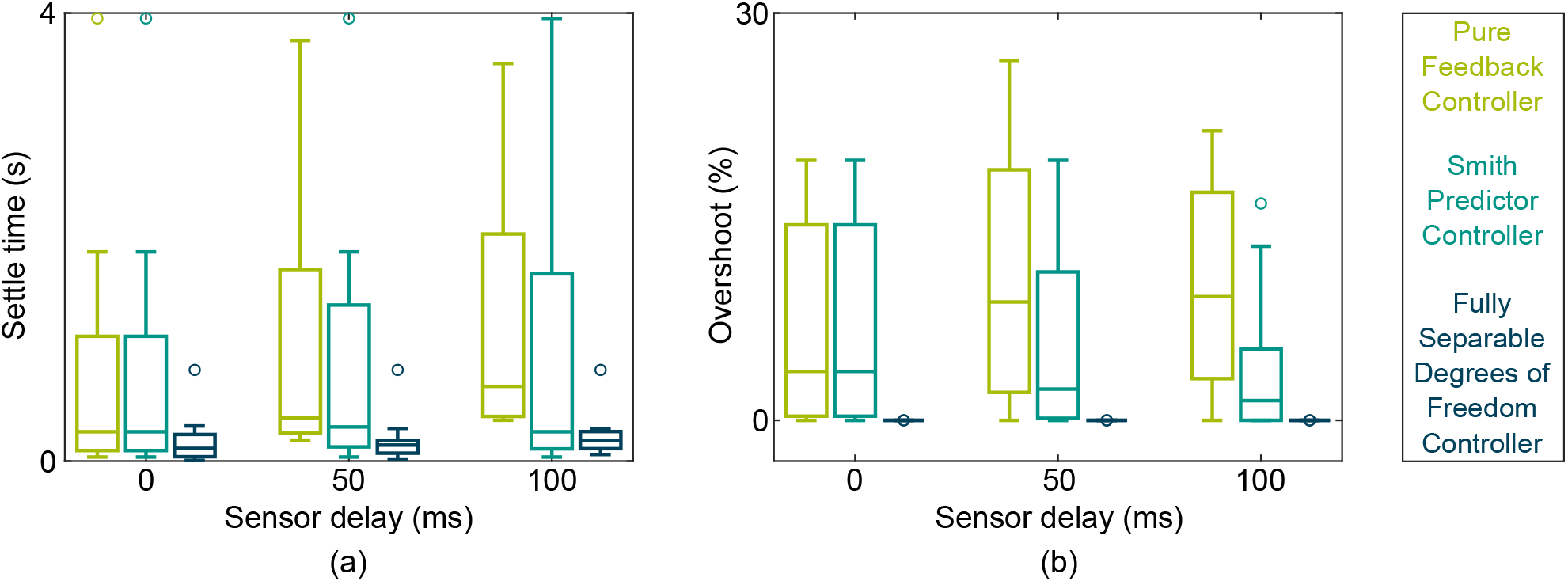
Performance of three different controllers with different sensor delays. (a) Settle time in seconds for each controller at different sensor delays. (b) Percentage overshoot of each controller at different sensor delays.

Whilst the FSDoF does not overshoot the desired state, for non-minimum phase systems i.e. plant structure 6 in Table 1, all controllers caused the state to initially go below zero before approaching the reference. In all cases, the FSDoF undershot by more than the SP and PFB for a similar maximum actuation. By adjusting FSDoF parameterization the undershoot can be arbitrarily reduced at the expense of settle time (Supp. Sec. 3).

The results from the Smith predictor (SP) should in theory be identical for all sensor delays. This is not the case for the results shown in Figure 3, as three cases were excluded from the analysis due to the PFB being unstable. For plant structure five, three parametrisations were unstable: *ω* = 10 rad s^*−*1^ *τ* = 100 ms, *ω* = 50 rad s^*−*1^ *τ* = 50 ms, and *ω* = 50 rad s^*−*1^ *τ* = 100 ms. When these are included in the analysis the Smith predictor has identical performance for all sensor delays - in line with theory.

A similar argument can be applied to the FSDoF controller. As seen in equation 7 the transfer function from the desired state to the actual state is independent of the sensor properties. This means the reference tracking performance is independent of any delays in the sensors, yet we see small variations in settle time as the sensor delay is increased. The reason for this variation is simply that due to the changing actuation required by the PFB, the FSDoF is allowed more or less actuation at different sensor delays. If the maximum actuation was fixed for all time delays, we would see the settle time being the same for different sensor delays, in line with the theory.

The settle time of the PFB increases significantly as the sensor delay increases. It goes from a mean of 0.79 s to 1.00 s to 1.18 s as the sensor delay increases. The Smith predictor has a mean settle time of 0.84 s which is reasonably consistent across different sensor delays. Both of these are much higher than the mean settle time of the FSDoF controller which has a mean settle time of 0.20 s; less than a quarter of the other two controllers. By all measures, the FSDoF is better than the PFB controller, and the performance gap increases with sensor delay. Even compared to a controller specifically designed to handle time delay systems, the SP, the FSDoF offers a large improvement. The handling of sensor delays is an inherent advantage of the FSDoF controller. It is worth noting that a feedback component is required in all three control architectures for rejecting disturbances, the disturbance rejection performance will suffer from increased sensor delays for all controllers.

### 4.2 Advantage 2: Mitigation of sensor noise

In classical feedback control, there is a trade-off between noise rejection and reference tracking. If the sensor signal is heavily filtered, the feedback control will be slowed down. If the feedback controller has a high bandwidth, it will track the desired state faster. However, it will come at the cost of a stronger response to noise from the sensors. A more formal mathematical evaluation of this principle is given in [25].

The behaviour of two PFB controllers highlights the trade-off. One of them rises to the desired state (unity) very quickly but is highly responsive to noise. The other one rejects the noise, however, it rises to the desired state much slower. The FSDoF control architecture rejects noise in the inner-loop, and changes state quickly with the noise-independent outer-loop.

With the fully-separable-degrees-of-freedom (FSDoF) control architecture, this trade-off exists for the inner-loop only. The outer-loop is not affected by sensor noise. As a result, and because the outer-loop controls all desired changes in state, the performance of reference tracking is independent of sensor noise. This is demonstrated in Figure 4.

**Figure 4:**
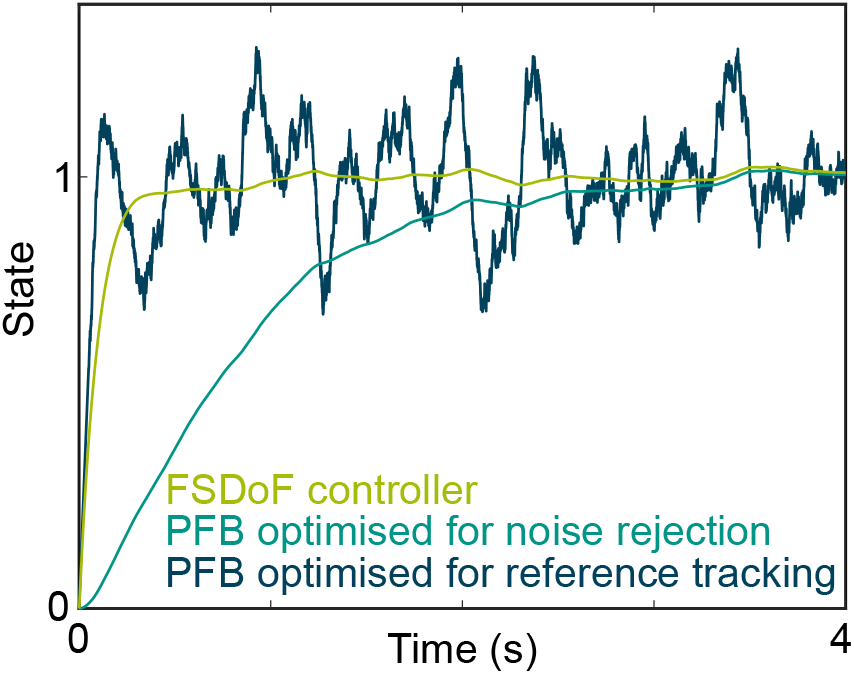
Example of the impact of sensor noise on reference tracking performance for different controllers: FSDoF, and two PFBs with different bandwidths.

For the Smith predictor (SP), a low-pass filter can be used to mitigate the impact of noise. This does not slow the response to desired changes since the feedback commands to change state are largely driven by the undelayed model, which does not contain noise. The low-pass filter is shown in Figure 1b as an optional filter. So, this advantage is apparent in both the SP and the FSDoF architectures.

### 4.3 Advantage 3: Preventing sensor saturation

In their study on efference copies in fruitflies, Kim et al. (2015) found the cancellation of the expected sensor response during spontaneous yaw turns to take place in specific directional-selective interneurons of the animal’s motion vision pathway [11]. Those so-called HS-cells, specifically the HSN- and HSE-cells, have been suggested to sense yaw rotations by processing wide-field optic flow [26]. Like any neuron in the nervous system, HS-cells have a limited electrical signalling range with which they encode motion information. By externally varying the membrane potential, the range of membrane currents of the HS-cells has been measured to cover approximately −25 to 10 nA [27]. Where the bounds are determined by the equilibrium potentials of contributing ion types. Cancelling the predicted sensory response of desired state changes, with an efference copy, keeps the signal of HS-cells within its operating range. Thereby, retaining high sensitivity to external disturbances, and effectively preventing saturation.

This is also the case for the output signals of spiking descending neurons which mediate sensory signals from the motion vision pathway and other modalities to the motor centres in the fly thoracic ganglion. Equation 8 is an *integrate and fire* model that can be used to describe the spike rate of a descending neuron. Where, *I* is the input current, *C* is the membrane capacitance, *V*_*th*_ is the threshold voltage, and *t*_*f*_ is a characteristic time for the neuron.

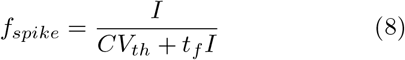

Clearly from this equation, if 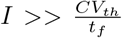 the neuron begins to saturate. Large changes in input current, result in small changes in spike rate.

If a fly did not cancel the expected sensor response during an intended bank turn, the visual neurons would be close to saturation and become less sensitive to external disturbances during the turn.

### 4.4 Controller robustness

Integrating all physiological and biomechanical properties of a biological control system would require an exceptional level of complexity. It is therefore reasonable to consider what happens when a controller is built around an incorrect or approximate model of the sensors, actuators and system dynamics.

#### 4.4.1 System dynamics modelling error

Firstly, we consider the case where the controller has been designed around system dynamics that are inaccurate or have changed. To address this we use a common metric in robust control, which we refer to as “stability robustness”. Stability robustness is a mathematical formulation for *how wrong the expected system dynamics can be such that the whole system retains stability*. [28]

Mathematically, we can answer this question by apsplying a multiplicative error to the system dynamics and deriving conditions for stability for the different controllers using the small-gain theorem [28]. Despite the desired change controller and forward model, the PFB and the FSDoF have the same stability robustness (see Appendix A) to “modelling” errors. This can be understood by considering the inner-loop of the FS-DoF controller as a mechanism that uses feedback to mitigate any unanticipated sensor response. This includes modelling errors: if the system dynamics differ from those expected by the forward model, the feedback controller will act to obtain the expected sensor response.

The stability robustness of the Smith predictor is not as simple. The small-gain theorem gives an identical result for the stability robustness for the same feedback controller *C*_1_(*s*). However, in the SP the feedback controller is not typically designed around controlling the delayed system, but rather the undelayed system, so *C*_1_(*s*) will not be the same. It is generally accepted that the Smith predictor is less robust than a simple feedback controller [29, 30], and some argue that for a fixed robustness a PID offers better performance than the SP (see [31]). The better performance by the SP over the PFB seen in this study, could just be a result of significantly lower stability margins, and therefore robustness.

#### 4.4.2 Sensor dynamics modelling error

An interesting feature of the FSDoF architecture is that instead of stabilising state units, it stabilises sensory units. For example in the fly visual system, instead of stabilising roll angle, a possible internal representation of the state, the inner loop would be stabilising the response of a neuron that is tuned to sense a roll rotation. This has some interesting effects: if the sensors act as high-pass filters, or derivatives, for the state units this could result in steady-state errors. This can be explained intuitively; the stabilising inner loop may not see the unexpected different sensor response for long enough to be able to mitigate it. This may appear to be a disadvantage, however, it works as an advantage for a different type of error. In biological systems, sensors are less reliable. Turning to the fly visual system again, the sensor response is dependent on the temporal and spatial properties of the environment [32]. If instead of the system dynamics being different to the expectation, the sensor response is: the resulting error due to this could be reduced.

In the two typical engineering controllers, if the sensors are incorrect, the feedback controller will settle to obtain an incorrect sensor response, thus the system will be at the wrong setpoint. With the FSDoF controller and a transient sensor response, a trade-off can be made. By permitting errors resulting from incorrect system dynamics, the errors from incorrect sensors signals will be reduced. It is worth noting that this trade-off can also impact the response to external disturbances. This effect is specific and depends on what system dynamics are controlled as well as the the properties of the sensors used.

To summarise, the risk of instability due to unexpected system dynamics for the FSDoF control architecture is identical to the PFB and SP controller. Secondly, in the PFB and SP, a system dynamics modelling error will generally not result in any steady state error between the actual state and the desired state. However, if there is an error in the sensor model they will stabilise around incorrect sensor values. The same applies to the FSDoF architecture when the sensors measure the absolute value of a given state variable. However, commonly biological sensors act like highpass filters or derivatives. In this case, with the FSDoF architecture, small steady-state errors may be permitted when the system dynamics are different to the expectation. The advantage of permitting that steady-state error is a reduced steady-state error when the sensor signal is incorrect.

## 5 Controller variants

There are demonstrable advantages of the FSDoF controller over a PFB. However, since the FSDoF is a form of the two-degree-of-freedom controller, there are equivalent variations shown in Figure 5. All five of those variants have the same transfer function as the FSDoF given by Equation 5, and therefore, in theory, identical reference tracking performance. The first three (Fig. 5a, 5b, 5c) have different structures to the FSDoF, using either a feedforward controller (Fig. 5a and 5c), or a reference filter alone (Fig. 5b). We will refer to these three variants as structurally different variants. The final two equivalent variations (Fig. 5d and 5e) use both a reference filter, and feedforward controller, akin to the FSDoF in Figure 1c, although in a different order. We will refer to these two variants as structurally similar variants. Experimental evidence suggests that biological control architectures send a feedforward command and cancels the expected sensor response [11, 12, 5]. This rules out the structurally different variants in Figure 5. However, there is no biological evidence in support of the original variant in Figure 1c, which does not also support the two structurally similar variants in Figure 5. The purpose of this section is to (i) explain the intuition behind these variants, (ii) explore reasons why biology might not use the structurally different variants, and (iii) assess practical differences of the structurally similar variants.

**Figure 5:**
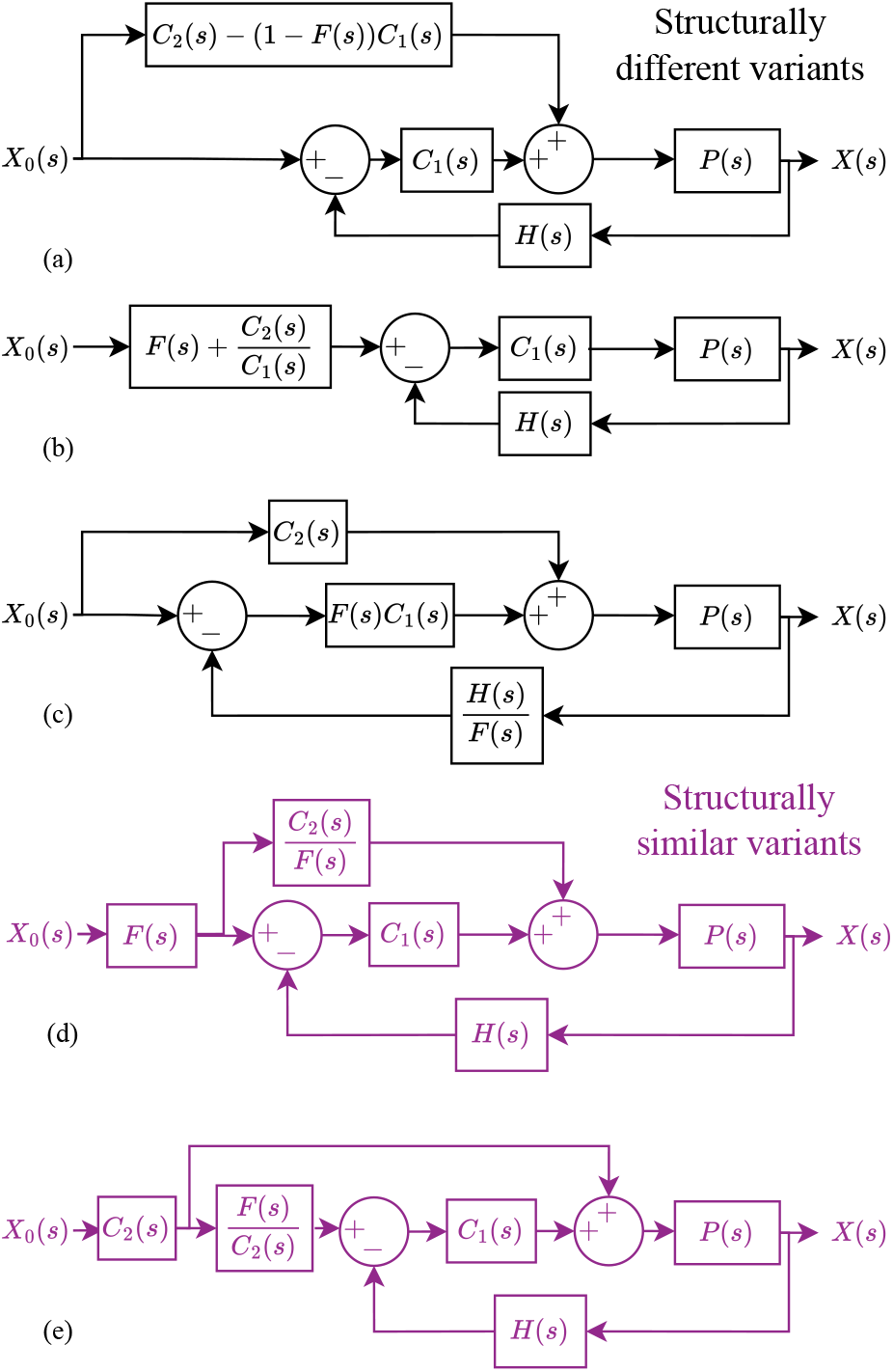
Five control architectures that give identical reference tracking equations to the FSDoF. Structurally different variants are given in black, and structurally similar variants in purple. We label the variants structurally similar if they are compatible with the electrophysiology results in [11, 12, 5]. (a) a feedforward motor command only. (b) a filter on the desired-state only. (c) The forward model is inside the feedback loop. (d) and (e) are equivalent structures to Figure 1c, both send the full actuation command in feedforward, and have forward models cancelling the expected sensor signal, just in an altered order.

### 5.0.1 Qualitative descriptions of the variants operation

The original structure in Figure 1c has two separate pathways from the desired state, one sending the actuation command, and one cancelling the expected sensor signal. In the first variant in Figure 5a the feedforward controller uses the forward model to provide the actuation required for the desired state change, minus, the actuation that the feedback controller is expected to generate based on the resulting sensor output. In this case, the cancellation occurs at the level of the actuator as opposed to the sensor. For example, if a fly intends to turn 10 rad s^*−*1^ to the left, it would “know” what motor command must be sent to the flight motor to produce the required torque. When the reafferent visual feedback triggered an optomotor response, it would also send a feedforward command to the muscles that inhibited that, i.e. cancel the expected motor command instead of the expected sensor response.

The variation in Figure 5b operates in the opposite way, it has no feedforward controller, just a filter on the reference. This reference filter cancels (i) the expected sensor response as a result of the desired state change, and (ii) the sensor response required to trigger the feedback controller to generate the actuation command. So, if a fly wanted to turn left, it would internally generate a sensor output that would trigger the optomotor left turn and then cancel the reafferent visual feedback.

Figure 5c shows a variant where the desired change controller has the same role, but the forward model has been moved inside the control loop. The sensor response is transformed into an estimation of the state by dividing by the forward model. compared to the desired state, and then feedback control is used.

Figure 5d shows a structurally similar variant where the forward model still cancels the expected sensor signal, and the feedforward controller provides all the actuation commands to change state. However, this time, the feedforward controller generates the actuation command based on the expected sensor response. If a fly wanted to turn left, it would send a cancellation of the expected sensor response - send the efference copy. Then, based on the efference copy signal it would generate the motor command to get that sensor response. The second structurally similar variant in Figure 5e operates the other way round. It generates the actuation command, and the forward model predicts the sensor response based on the actuation command. This final variant reflects more directly the original etymology of efference copy, meaning motor copy. A fly turning left would send the motor command to the actuator, and then, based on the motor command signal it would generate the expected sensor response it should cancel.

### 5.0.2 Why not cancel the expected motor command?

Flies appear to cancel the expected sensor signal as opposed to the motor command initiated by the reafferent sensor response as suggested in Figure 5a. The first reason may be that this variant does not always exhibit internal stability, whilst it is input-to-output stable, there are conditions under which individual blocks are not. These scenarios arise when *C*_1_ has an integral component. When there is integral action any combination of input signal (*X*_0_) and forward model (*F*) such that (1 *− F*)*X*_0_ has a non-zero average, will lead to internal instability. The integral components in the feed-forward and feedback paths will tend towards *±∞*. A non-zero average in (1 *− F*)*X*_0_ can arise when (i) there are high-pass filter sensors, (ii) there is a non-linearity, for example, if there is a limit on the rate at which *F*(*s*) can change its output, or (iii) if the implemented *F*(*s*) does not have a perfect unity DC gain.

The variant in Figure 5c also has a potential problem, in this case with the inversion of the forward model *F*(*s*). Firstly, if there is a positive zero in the system dynamics, a positive zero in *F*(*s*) is required for *F*(*s*) = *C*_2_*PH* to hold. An inversion of *F*(*s*) under this condition, as in Fig. 5c, will result in an unstable pole. Secondly, if there is a time delay, the inversion will create the need for a positive time delay which is not possible.

Lastly, and perhaps most significantly, none of the three structurally different architectures prevents sensor output saturation, as described in Section 4.3.

### 5.0.3 Does the order matter?

Considering the two structurally similar variants, they both solve the sensor saturation problem. The first of the two variants generates the feedforward motor command based on the efference copy. This variant suffers the same challenges with inverting *F*(*s*) as explained above. If the plant dynamics had a positive zero, the inversion of this would lead to breaking internal stability.

The final structurally similar variant (Fig. 5e) is more plausible and reflects the etymological origins of how efference copies may be used in sensorimotor control. One practical challenge with this architecture however is its handling of unstable plant dynamics. Unstable plant dynamics have a positive pole in *P*(*s*). The filter 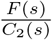, is a model of the plant and sensor dynamics, from Equation 6. Is it possible for animals to have a neural filter that represents unstable dynamics?

In the structure analysed in Figure 1c each block in the diagram is causal. In addition, when controlling unstable dynamics and a stabilising desired change controller, the forward model will be internally stable. It is worth noting that there have not been physiological experiments showing the exact origin of the feedforward control command generated by the desired change controller *C*_2_(*s*), or the efference copy generated by the forward model *F*(*s*). In stable biological systems, it is perfectly possible to have a structure similar to that shown in Figure 5e.

## 6 Conclusions

Motivated by recent studies in the insect sensorimotor system we developed a phenomenological control architecture and benchmarked its performance against common control engineering approaches. We find that for accurate fully-separable-degrees-of-freedom (FSDoF) controllers, the performance in tracking the desired state is independent of the sensor output. This means that the control architecture is particularly good at handling sensors with time delays and significant noise.

Sensory delays are common in biological systems [33, 34, 35], yet many animals employ high-performance control systems. Our work suggests how an FSDoF controller with forward models (efference copies) and desired change controllers could achieve this. In particular for systems with fast dynamics but significant sensor delays, like insects. This combination of fast dynamics and comparatively slow sensors is present in the fly visual system and could be a reason why efference copies were first recorded there [35, 32].

Another factor to consider is noise in the nervous system [36]. We show that the FSDoF architecture allows fast goal-directed behaviours despite sensor noise. This is due to the inherent property of the outer loop control signals being sensor-independent. Secondly, by subtracting the expected sensor response the sensors are kept in their operating range, maintaining sensitivity to external perturbations. This could cause a stronger signal power, and therefore, for a fixed level of noise, a better signal-to-noise ratio.

We show that the FSDoF control architecture does not overshoot, which is consistent with experiments where humans are tasked with reaching an object and the position of their hand is tracked [37, 21].

Whilst the PFB makes up 90 to 95% of industrial control methods [38], there are other architectures we have not compared here. One such architecture is the model predictive controller (MPC), which each time step optimises future actuation commands to minimise a balance of actuation and error [39]. This has significantly more computational complexity, and it is unlikely that biology could implement this kind of structure.

The visual system in flies, where there is physiological evidence for efference copies, does not respond linearly to velocity. Instead, it has a response that contains significant modulations that depend on the spatio-temporal frequency distribution of the environment [40, 35]. Despite this, flies remain incredibly agile and fast. We have explained the ability of the FSDoF architecture to suppress sensor noise while enabling fast desired state changes. This could explain how the insect flight control system effectively operates despite these modulations from the visual sensors.

A point that raises interesting questions is that the feedback controller does not control parameters in state units, but instead controls parameters in sensory units. This could potentially lend itself to a distributed control system. Instead of having many sensors integrated to provide the current state and then having a single, complicated, feedback controller. It allows each sensor system to sense something unanticipated and feedback directly to a corresponding motor system. This could potentially reduce computational overheads, inverting and combining information from multiple sensors, and provide a shorter neural pathway from sensor to muscle. Thereby, reducing feedback delays, and improving disturbance rejection performance. There is another potential advantage of this strategy. Different sensing modalities have different response delays. The FSDoF control architecture could provide a mechanism to utilise sensors operating at different speeds. A similar strategy was found in the fly gaze stabilisation system where the fast mechanoreceptor signals provide muscle commands quickly, while the slower visual system incurs a longer response delay before activating the neck muscles [41].

Another quality of the FSDoF controller is that the desired changes in state, when given unlimited actuation capabilities, can be arbitrarily fast (Supp. Sec. 4). This is not the case for the SP or PFB where high gains can lead to instability, in particular for systems with sensor delays. The only trade-off with reducing the settle time of the FSDoF, except for the increased actuation, is for plant 6 where the undershoot significantly increases. Further biological system identification [42] studies would shed more light on whether non-minimum phase systems are present in insect dynamics.

Finally, there is a distinction made in recent work between “graded” and “all-or-none” efference copies [14]; where a graded efference copy is the output of a forward model predicting the sensor response to cancel, and all-or-none is effectively turning off the sensor when enacting a desired state change. In this paper we have analysed a graded efference copy model, however, animals likely employ both methods. The results here can be extrapolated to all-or-none efference copy control systems; the three advantages would still be obtained. The two primary differences in performance are that with the all-or-none efference copy (i) it becomes theoretically impossible to control unstable systems, and (ii) external disturbances during desired state changes are not sensed.

This paper demonstrates potential advantages to biological systems using control structures like the efference copy controller described here. In particular given the agility and high-speed manoeuvres required in nature, with the differently delayed sensor signals. Future work should address how these advantages can be retained through changing system dynamics. Whilst we show that the FSDoF is equally robust as the PFB, the performance is better when the controller is accurate. Further work on adaptation in the insect sensorimotor system could provide valuable insights into how this is achieved, both in biological sensors [43, 44, 45] and during different locomotor states [46, 47, 48, 49].

To summarise, we explore a control architecture based on principles found in insect sensorimotor control. We show the FSDoF architecture has three advantages: (i) it handles systems with time delays better than the PFB and SP, (ii) it is an inherently noise-robust architecture, and (iii) it provides a natural mechanism for keeping sensors in their operating range. These findings point towards the suitability of such architectures in biology and aim to show the advantages that efference copies can have in sensorimotor control.

## A Appendix 1

First, we look at the robustness of the control schemes to system dynamics that are different to the expectation. A multiplicative error can be modelled in the Laplace domain by making the plant *P*(*s*)(1 + Δ_*P*_ (*s*)); where Δ_*P*_ (*s*) is the Laplace transform of the error, and *P*(*s*) is the model of the plant dynamics that were expected [28]. The conditions for closed-loop stability for all three controllers can be derived as an extension to Nyquist using the small-gain theorem [28].

### A.0.1 FSDoF and PFB Robustness

The result of a sufficient condition for closed-loop stability is given in Equation 9

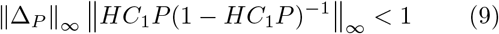

If this condition is satisfied, the system will be closed-loop stable [28]. It is immediate that the transfer functions *C*_2_(*s*), and *F*(*s*) do not appear in Condition 9, and if they are internally stable, they will not cause instability.

Even if the system with unexpected system dynamics remains stable, the performance can still suffer. To evaluate this we can find the transfer functions for the three control architectures including the multiplicative error. First, the FSDoF controller we substitute the plant with error into Equation 5, and leave the forward model unchanged. That gives the transfer function in Equation 10.

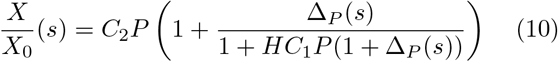

For the PFB the transfer function is given in equation 11.

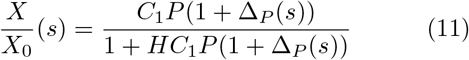

### A.0.2 SP Robustness

Following the notation in Figure 1, *P*(*s*)*H*(*s*) = *G*(*s*)*e*^*−τs*^, the SP feedback controller is designed around *G*(*s*). When the multiplicative modelling error is included in the transfer function the original cancellation no longer occurs as shown in equation 12.

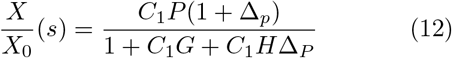

However, since the response to disturbances is the same, the small-gain theorem gives the same result.

