## Supplementary for "A control engineering perspective on the advantages of efference copies"

### Supplementary information: A control engineering perspective on the advantages of efference copies

May 15, 2024

Imperial College London, Exhibition Rd, South Kensington, London, SW7 2BX

#### 1 Results

Table 1: Pure feedback controller (PFB) table of results

| Plant structure | $\omega$ (rad/s) | $\tau$ (s) | Max actuation | Total actuation | Settle time (s) | Overshoot (%) |
|---|---|---|---|---|---|---|
| 1 | 10 | 0 | 1.7212 | 0.22457 | 0.1784 | 0.61067 |
| 1 | 10 | 0.1 | 1.4257 | 0.58452 | 0.56067 | 8.25 |
| 1 | 10 | 0.05 | 1.8365 | 0.38207 | 0.32206 | 8.7177 |
| 1 | 50 | 0 | 1.7212 | 0.044918 | 0.035685 | 0.61067 |
| 1 | 50 | 0.1 | 1.1479 | 0.38212 | 0.39842 | 13.2661 |
| 1 | 50 | 0.05 | 1.1483 | 0.22819 | 0.23443 | 10.716 |
| 2 | 10 | 0 | 199.2938 | 2.529 | 0.3046 | 2.3031 |
| 2 | 10 | 0.1 | 50.9481 | 1.3599 | 0.63634 | 8.8128 |
| 2 | 10 | 0.05 | 114.9587 | 2.242 | 0.25202 | 8.3084 |
| 2 | 50 | 0 | 333.6711 | 3.1107 | 0.065618 | 15.9754 |
| 2 | 50 | 0.1 | 40.5438 | 1.5825 | 0.69682 | 21.328 |
| 2 | 50 | 0.05 | 59.7381 | 1.8048 | 0.38318 | 21.039 |
| 3 | 10 | 0 | 3.4534 | 0.65981 | 0.55488 | 4.902 |
| 3 | 10 | 0.1 | 9.9907 | 0.8442 | 0.70429 | 18.0198 |
| 3 | 10 | 0.05 | 12.9328 | 0.53905 | 0.39502 | 9.2305 |
| 3 | 50 | 0 | 2.2945 | 0.12968 | 0.10865 | 5.2519 |
| 3 | 50 | 0.1 | 1.1253 | 0.38351 | 0.39139 | 9.4171 |
| 3 | 50 | 0.05 | 1.1381 | 0.1905 | 0.18627 | 7.6357 |
| 4 | 10 | 0 | 1.6443 | 0.27218 | 0.21915 | 12.8313 |
| 4 | 10 | 0.1 | 1.7413 | 0.41552 | 0.36451 | 16.7991 |
| 4 | 10 | 0.05 | 2.3466 | 0.32315 | 0.25 | 26.516 |
| 4 | 50 | 0 | 0.99995 | 2.9595 | 3.9527 | -0.0051387 |
| 4 | 50 | 0.1 | 0.99998 | 2.6544 | 3.5496 | -0.0016679 |
| 4 | 50 | 0.05 | 0.99997 | 2.8112 | 3.7553 | -0.0030613 |
| 5 | 10 | 0 | 8.4098 | 0.47791 | 0.39104 | 19.1577 |
| 5 | 10 | 0.05 | 13.2699 | 2.7653 | 2.0451 | 157.6398 |
| 5 | 50 | 0 | 8.4098 | 0.095586 | 0.078212 | 19.1577 |
| 6 | 10 | 0 | 1 | 1.386 | 1.8682 | 0.0014718 |
| 6 | 10 | 0.1 | 1.0269 | 1.9828 | 2.4505 | 3.0725 |
| 6 | 10 | 0.05 | 1.0018 | 1.3164 | 1.8371 | 0.20327 |
| 6 | 50 | 0 | 1 | 1.2987 | 1.6725 | -4.0536e-10 |
| 6 | 50 | 0.1 | 1 | 1.4947 | 2.028 | -4.3091e-09 |
| 6 | 50 | 0.05 | 1 | 0.9747 | 1.3301 | -3.6593e-10 |

Table 2: Smith predictor (SP) table of results

| Plant structure | $\omega$ (rad/s) | $\tau$ (s) | Max actuation | Total actuation | Settle time (s) | Overshoot (%) |
|---|---|---|---|---|---|---|
| 1 | 10 | 0 | 1.7212 | 0.22457 | 0.1784 | 0.61067 |
| 1 | 10 | 0.1 | 1.7212 | 0.22457 | 0.1784 | 0.61067 |
| 1 | 10 | 0.05 | 1.7212 | 0.22457 | 0.1784 | 0.61067 |
| 1 | 50 | 0 | 1.7212 | 0.044918 | 0.035685 | 0.61067 |
| 1 | 50 | 0.1 | 1.7212 | 0.044918 | 0.035685 | 0.61067 |
| 1 | 50 | 0.05 | 1.7212 | 0.044918 | 0.035685 | 0.61067 |
| 2 | 10 | 0 | 199.2938 | 2.529 | 0.3046 | 2.3031 |
| 2 | 10 | 0.1 | 199.2938 | 2.529 | 0.3046 | 2.3031 |
| 2 | 10 | 0.05 | 199.2938 | 2.529 | 0.3046 | 2.3031 |
| 2 | 50 | 0 | 333.6711 | 3.1107 | 0.065619 | 15.9754 |
| 2 | 50 | 0.1 | 333.6711 | 3.1107 | 0.065619 | 15.9754 |
| 2 | 50 | 0.05 | 333.6711 | 3.1107 | 0.065619 | 15.9754 |
| 3 | 10 | 0 | 3.4534 | 0.65981 | 0.55488 | 4.902 |
| 3 | 10 | 0.1 | 3.4534 | 0.65981 | 0.55488 | 4.902 |
| 3 | 10 | 0.05 | 3.4534 | 0.65981 | 0.55488 | 4.902 |
| 3 | 50 | 0 | 2.2945 | 0.12968 | 0.10865 | 5.2519 |
| 3 | 50 | 0.1 | 2.2945 | 0.12968 | 0.10865 | 5.2519 |
| 3 | 50 | 0.05 | 2.2945 | 0.12968 | 0.10865 | 5.2519 |
| 4 | 10 | 0 | 1.6443 | 0.27218 | 0.21915 | 12.8313 |
| 4 | 10 | 0.1 | 1.6443 | 0.27218 | 0.21915 | 12.8313 |
| 4 | 10 | 0.05 | 1.6443 | 0.27218 | 0.21915 | 12.8313 |
| 4 | 50 | 0 | 0.99995 | 2.9595 | 3.9527 | -0.0051387 |
| 4 | 50 | 0.1 | 0.99995 | 2.9595 | 3.9527 | -0.0051387 |
| 4 | 50 | 0.05 | 0.99995 | 2.9595 | 3.9527 | -0.0051387 |
| 5 | 10 | 0 | 8.4098 | 0.47791 | 0.39104 | 19.1577 |
| 5 | 10 | 0.05 | 8.4098 | 0.47791 | 0.39104 | 19.1577 |
| 5 | 50 | 0 | 8.4098 | 0.095586 | 0.078212 | 19.1577 |
| 6 | 10 | 0 | 1 | 1.386 | 1.8682 | 0.0014718 |
| 6 | 10 | 0.1 | 1 | 1.386 | 1.8682 | 0.0014718 |
| 6 | 10 | 0.05 | 1 | 1.386 | 1.8682 | 0.0014718 |
| 6 | 50 | 0 | 1 | 1.2987 | 1.6725 | -4.0536e-10 |
| 6 | 50 | 0.1 | 1 | 1.2987 | 1.6725 | -4.0536e-10 |
| 6 | 50 | 0.05 | 1 | 1.2987 | 1.6725 | -4.0536e-10 |

Table 3: Fully-seperable-degrees-of-freedom (FSDoF) controller table of results

| Plant structure | $\omega$ (rad/s) | $\tau$ (s) | Max actuation | Total actuation | Settle time (s) | Overshoot (%) |
|---|---|---|---|---|---|---|
| 1 | 10 | 0 | 1.708 | 0.22461 | 0.15501 | -1.5787e-11 |
| 1 | 10 | 0.1 | 1.4121 | 0.59165 | 0.20134 | -2.2826e-11 |
| 1 | 10 | 0.05 | 1.8227 | 0.37316 | 0.14262 | -1.5743e-11 |
| 1 | 50 | 0 | 1.7055 | 0.044904 | 0.031065 | -3.9191e-12 |
| 1 | 50 | 0.1 | 1.1379 | 0.39851 | 0.05807 | -7.816e-12 |
| 1 | 50 | 0.05 | 1.1384 | 0.23454 | 0.058016 | -7.7827e-12 |
| 2 | 10 | 0 | 199.3839 | 2.7954 | 0.13558 | -1.6875e-11 |
| 2 | 10 | 0.1 | 50.6912 | 1.3929 | 0.26331 | -2.7456e-11 |
| 2 | 10 | 0.05 | 114.3585 | 2.0797 | 0.17768 | -1.3689e-11 |
| 2 | 50 | 0 | 335.1802 | 1.5862 | 0.046018 | -5.9841e-12 |
| 2 | 50 | 0.1 | 40.3913 | 1 | 0.12982 | -1.7808e-11 |
| 2 | 50 | 0.05 | 60.0318 | 1 | 0.10649 | -1.2257e-11 |
| 3 | 10 | 0 | 3.4643 | 0.64743 | 0.31345 | -3.4639e-11 |
| 3 | 10 | 0.1 | 9.9555 | 0.8409 | 0.1849 | -2.2804e-11 |
| 3 | 10 | 0.05 | 12.8506 | 0.53923 | 0.16275 | -1.7275e-11 |
| 3 | 50 | 0 | 2.2862 | 0.1222 | 0.077169 | -7.1942e-12 |
| 3 | 50 | 0.1 | 1.1174 | 0.39355 | 0.11039 | -1.0014e-11 |
| 3 | 50 | 0.05 | 1.1296 | 0.18863 | 0.10979 | -9.9698e-12 |
| 4 | 10 | 0 | 1.6288 | 0.25412 | 0.091489 | 2.2427e-12 |
| 4 | 10 | 0.1 | 1.7247 | 0.40496 | 0.076459 | 5.5067e-12 |
| 4 | 10 | 0.05 | 2.3247 | 0.30905 | 0.022104 | 2.065e-12 |
| 4 | 50 | 0 | 1 | 3.8393 | 0.2917 | -1.3323e-12 |
| 4 | 50 | 0.1 | 1 | 3.4363 | 0.2917 | -1.3323e-12 |
| 4 | 50 | 0.05 | 1 | 3.642 | 0.2917 | -1.3323e-12 |
| 5 | 10 | 0 | 8.3439 | 0.44429 | 0.028242 | -2.8422e-12 |
| 5 | 10 | 0.05 | 13.1447 | 2.1131 | 0.017309 | -2.3537e-12 |
| 5 | 50 | 0 | 8.3347 | 0.088853 | 0.0056623 | -1.2212e-13 |
| 6 | 10 | 0 | 1 | 1.746 | 0.81366 | -1.5312e-10 |
| 6 | 10 | 0.1 | 1 | 2.3283 | 0.81366 | -1.5312e-10 |
| 6 | 10 | 0.05 | 1 | 1.7149 | 0.81366 | -1.5312e-10 |
| 6 | 50 | 0 | 1 | 1.6516 | 0.18295 | -1.7124e-10 |
| 6 | 50 | 0.1 | 1 | 2.0072 | 0.18295 | -1.7124e-10 |
| 6 | 50 | 0.05 | 1 | 1.3093 | 0.18295 | -1.7124e-10 |

### 2 Controller structure and tuning

#### 2.1 FSDoF tuning

To allow for a fair comparison it was important to optimise $F(s)$ and $C_2(s)$ with a single degree of freedom, i.e. one value which we call $q$. Two things should be satisfied for a legitimate FSDoF controller in the context of these simulations:

$$F(s) = C_2PH(s) \tag{1}$$

$$\lim_{s\to 0} F(s) = 1 \tag{2}$$

$H(s) = 1$ in all the simulations, and assuming the plant dynamics were of the form:

$$P(s) = \frac{P_n(s)}{P_d(s)}e^{-\tau s} \tag{3}$$

We then take the roots of the numerator polynomial such that:

$$P_n(s) = P_n^s(s)P_n^u(s) \tag{4}$$

Where $P_n^s(s)$ and $P_n^u(s)$ are the polynomials of all the negative zeros and positive zeros of the numerator respectively. Substituting this provides:

$$F(s) = C_2\frac{P_n^s P_n^u}{P_d}e^{-\tau s} \tag{5}$$

By letting:

$$C_2(s) = \frac{P_d}{\left(\frac{s}{q} - 1\right)^2 P_n^s} \tag{6}$$

$$F(s) = \frac{P_n^u}{a\left(\frac{s}{q} - 1\right)^2} e^{-\tau s} \tag{7}$$

Where:

$$a = \lim_{s \to 0} P_n^u(s) \tag{8}$$

We can meet the two requirements in Equation 1 and 2, ensure $F(s)$ and $C_2(s)$ are internally stable, and most importantly the FSDoFs reference tracking performance is controlled by a single variable $q$.

#### 2.2 Controller values

Table 4: Table of controller parameterisations. The Smith predictor had the same gains as the PFB when $\tau = 0$ for all values of $\tau$.

| Plant structure | $\omega$ (rad/s) | $\tau$ (s) | PFB: K$_p$ | PFB: K$_i$ | PFB: K$_d$ | FSDoF: $q$ |
|---|---|---|---|---|---|---|
| 1 | 10 | 0 | 1.7212 | 19.5871 | 0 | 37.6369 |
| 1 | 10 | 0.1 | 0.86007 | 5.6566 | 0 | 28.9758 |
| 1 | 10 | 0.05 | 1.2761 | 11.2087 | 0 | 40.9076 |
| 1 | 50 | 0 | 1.7212 | 97.9357 | 0 | 187.825 |
| 1 | 50 | 0.1 | 0.40866 | 7.3928 | 0 | 100.4709 |
| 1 | 50 | 0.05 | 0.42975 | 13.1689 | 0 | 100.5642 |
| 2 | 10 | 0 | 21.2534 | 6.4618 | 1.6781 | 43.0316 |
| 2 | 10 | 0.1 | 5.0023 | 0.42403 | 0.45127 | 22.1563 |
| 2 | 10 | 0.05 | 9.2896 | 1.5187 | 0.99159 | 32.8344 |
| 2 | 50 | 0 | 106.2671 | 161.5462 | 1.6781 | 126.7761 |
| 2 | 50 | 0.1 | 7.3683 | 0.97547 | 0.3081 | 44.9396 |
| 2 | 50 | 0.05 | 12.8405 | 2.9942 | 0.41317 | 54.7867 |
| 3 | 10 | 0 | 1.9691 | 10.5361 | 0.014843 | 18.6125 |
| 3 | 10 | 0.1 | 1.4511 | 6.1648 | 0.085395 | 31.5524 |
| 3 | 10 | 0.05 | 2.1159 | 9.1138 | 0.10817 | 35.8477 |
| 3 | 50 | 0 | 1.9691 | 52.6805 | 0.0029686 | 75.6006 |
| 3 | 50 | 0.1 | 0.43646 | 6.7586 | 0.0051959 | 52.8529 |
| 3 | 50 | 0.05 | 0.51857 | 11.5895 | 0.0051033 | 53.141 |
| 4 | 10 | 0 | 0.16939 | 56.601 | 0 | 63.7718 |
| 4 | 10 | 0.1 | 0.8655 | 8.7584 | 0 | 76.3081 |
| 4 | 10 | 0.05 | 1.2446 | 22.0426 | 0 | 263.9988 |
| 4 | 50 | 0 | 0 | 0.99964 | 0 | 20 |
| 4 | 50 | 0.1 | 0 | 0.99964 | 0 | 20 |
| 4 | 50 | 0.05 | 0 | 0.99964 | 0 | 20 |
| 5 | 10 | 0 | -7.1377 | -40.482 | 0 | 206.5766 |
| 5 | 10 | 0.05 | -1.5703 | -2.2557 | -0.072806 | 337.1645 |
| 5 | 50 | 0 | -7.1377 | -202.4101 | 0 | 1031.6281 |
| 6 | 10 | 0 | 0 | 1.0905 | 0 | 9 |
| 6 | 10 | 0.1 | 0 | 1.0005 | 0 | 9 |
| 6 | 10 | 0.05 | 0 | 0.99738 | 0 | 9 |
| 6 | 50 | 0 | 0 | 1.3688 | 0 | 49 |
| 6 | 50 | 0.1 | 0 | 1.0093 | 0 | 49 |
| 6 | 50 | 0.05 | 0 | 1.3153 | 0 | 49 |

### 3 Plant structure 6 and non-minimum phase systems

For the non-minimum phase system in plant 6, $P_n^u$ is a single-order polynomial. This means there is an initial undershoot of 0 in the state before settling at the desired state. Figure 1 demonstrates that adjusting $q$ allows you to balance undershoot and settle time.

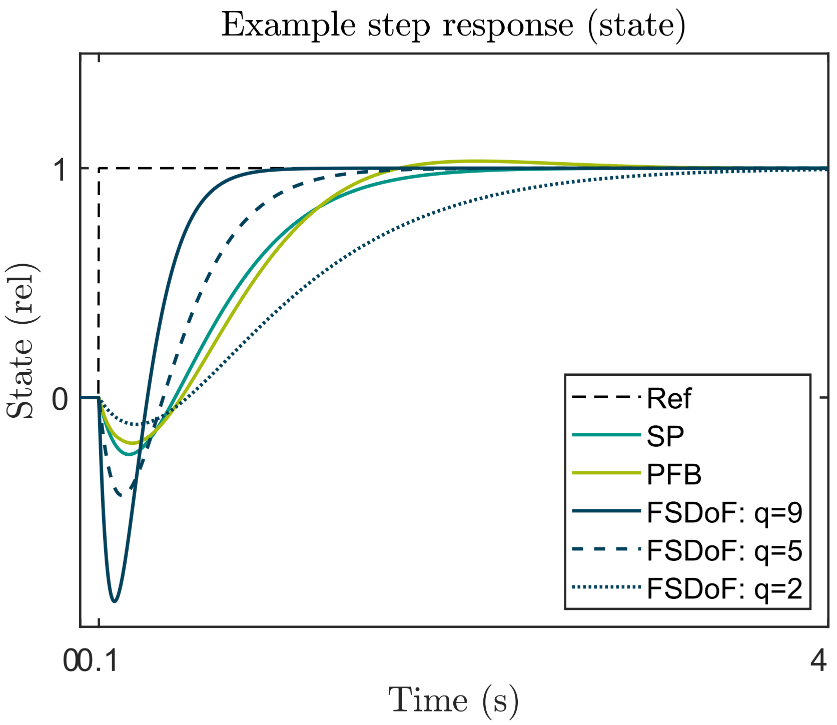


Figure 1: Step response of the SP, PFB, and three FSDoF controllers with different values of $q$. As $q$ is increased the undershoot increases, but the settle time reduces. This was plant structure 6 with $\omega = 10$ rad/s and $\tau = 100$ ms.

### 4 Arbitrarily fast settle time of the FSDoF

As explained in the manuscript, unlike the PFB and SP, the FSDoF can settle arbitrarily quickly by increasing $q$ when there is no limit on the maximum actuation available. Examples of this are shown in Figure 2, where the initial value of $q$ is shown, and a simulation with $q$ increased by a factor of 5 is plotted. The only time there is a significant trade-off, not counting increased actuation, is for plant number 6 as shown above where the undershoot dramatically increases.

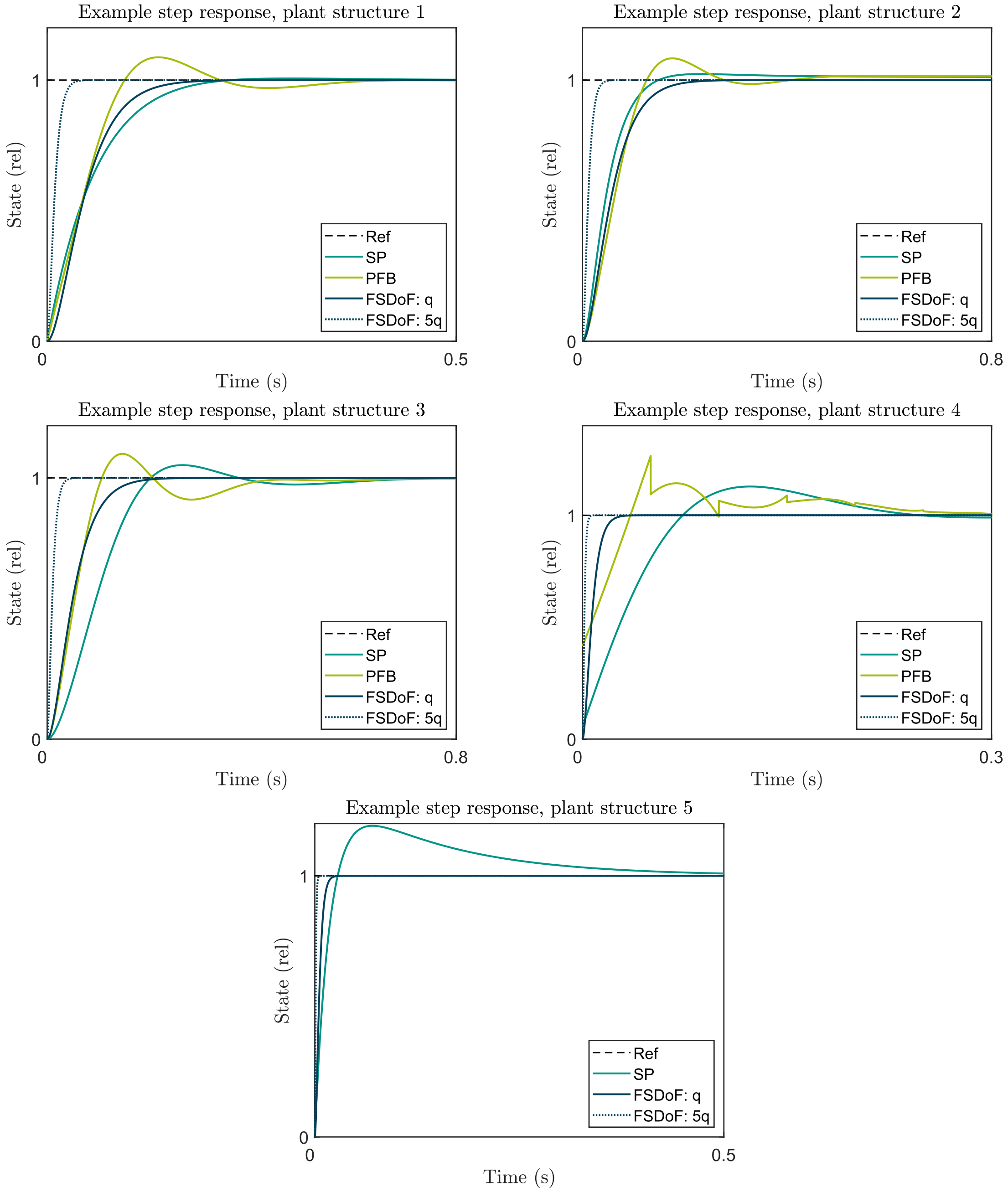


Figure 2: Demonstrating how $q$ can be arbitrarily increased for a faster settle time. Each plant structure is parameterised by $\tau = 50$ ms and $\omega = 10$ rad/s. The example for plant 5 does not include the PFB because it was significantly worse and made the FSDoF results difficult to see when scaled.
